# Behavioral pharmacology of mescaline - the role of serotonin 5-HT2A, 5-HT2B, 5-HT2C and 5-HT1A receptors

**DOI:** 10.1101/2024.08.28.610032

**Authors:** Lucie Olejníková-Ladislavová, Michaela Fujáková-Lipski, Klára Šíchová, Hynek Danda, Kateřina Syrová, Jiří Horáček, Tomáš Páleníček

## Abstract

**Rationale:** Mescaline is a classical psychedelic compound with a phenylethylamine structure that primarily acts on serotonin 5-HT2A/C receptors, but also binds to 5-HT1A and 5-HT2B receptors. Despite being the first psychedelic ever isolated and synthesized, the precise role of different serotonin receptor subtypes in its behavioral pharmacology is not fully understood.

**Objectives:** In this study, we aimed to investigate how selective antagonists of 5-HT2A, 5-HT2B, 5-HT2C, and 5-HT1A receptors affect the behavioral changes induced by subcutaneous administration of mescaline (at doses of 10, 20, and 100 mg/kg) in rats.

**Methods:** We used adult male Wistar rats in all our experiments. We evaluated locomotor activity using the open field test, and assessed sensorimotor gating deficits by measuring prepulse inhibition (PPI) of acoustic startle reaction (ASR).

**Results:** While the highest dose of mescaline induced hyperlocomotion, which almost all the other antagonists reversed, the PPI deficits were selectively normalized by the 5-HT2A antagonist. The 5-HT2C antagonist partially reversed the small decrease in locomotor activity induced by lower doses of mescaline.

**Conclusion:** Our findings suggest that mescaline-induced changes in behavior are primarily mediated by the 5-HT2A receptor subtype, with less pronounced contributions from the 5-HT2C receptor. The other antagonists had limited effects.

## 1 INTRODUCTION

Mescaline, a member of the phenylethylamine group of serotonergic hallucinogens, is found in various members of the Cactaceae family, with peyote (Lophophora williamsii) and San Pedro (Echinopsis pachanoi) cactus being the most well-known sources. It was first isolated by German scientist Arthur Carl Wilhelm Heffter in 1897, making it the first known psychedelic recognized by Western scientists (Handbook of Experimental Pharmacology).

Compared to other phenylethylamine and tryptamine hallucinogens, mescaline has relatively moderate affinities towards serotonin (5-hydroxytryptamine, 5-HT) receptors. This is consistent with the need for higher doses (hundreds of milligrams per dose) to achieve a full psychedelic effect in humans [1]. The affinities of mescaline have been reported to be in the following order: 5-HT1A > 5-HT2A > 5-HT2C > 5-HT2B, with a 5-HT2A: 5-HT1A ratio of 0.73 and a 5-HT2A: 5-HT2C ratio of 2.7. [2,3].

Rodent studies, including one from the authors’ laboratory, suggest that mescaline has a dose and time-dependent biphasic mode of action on locomotor behavior. The effects also depend on the rodent model used and the form of administration. However, consistently high doses of mescaline induce initial inhibition followed by an increase in locomotion, which seems to be linked to 5-HT2A agonism [4–6]. Other preclinical reports from the late 1970s indicate that high doses of mescaline can induce pathological aggression, which is not observed after other hallucinogenic phenylethylamines [7,8]. This biphasic mode of action raises the possibility of differential involvement of serotonin receptors in the pharmacodynamics of mescaline.

While the effects of serotonergic psychedelics in other behavioral paradigms, such as head-twitch response, and disruption of the prepulse inhibition (PPI), seem to be also 5-HT2A-mediated [4,5,9–11], the contribution of other 5-HT receptors is less clear. For example, previous studies with the tryptamine hallucinogen psilocin have shown that the locomotor inhibitory action of psilocin is 5-HT2C and 5-HT1A mediated and that 5-HT2A receptors do not contribute at all [12]. Similarly, phenylethylamine hallucinogens 2,5-dimethoxy-4-iodoamphetamine (DOI) and psilocin have shown that activation of the 5-HT2A receptor induces an increase in locomotor activity at low doses, while a decrease in locomotion at high doses is more likely associated with agonism of the 5-HT2C receptor [12–15]. Additionally, DOI-induced sensorimotor gating deficits were reversed by both a selective 5-HT2A antagonist [16], and a selective 5-HT2C receptor agonist [17]. The role of 5-HT1A receptors is also under-investigated; however, there is some evidence of their contribution to DOI-induced PPI deficits [18] or psilocin-induced hypolocomotion [12].

Main objective of the current study was to investigate the contribution of serotonin 5-HT1A, 5-HT2A, 5-HT2B, and 5-HT2C receptors to behavioral changes induced by mescaline using specific antagonists administration. The study focused on two behavioral paradigms: 1) mescaline’s biphasic effects on locomotion in the open field test (OFT) and 2) its disruptive effect on sensorimotor processing, specifically prepulse inhibition of acoustic startle reaction (PPI ASR). Selective antagonists of all the above-mentioned 5-HT receptors were used to study the pharmacodynamics of behavioral changes.

## 2 METHODS

This study represents a reanalysis and expansion of a previously published work of Paleníček et al. from 2008 [19], where the impact of three varying doses of mescaline (10, 20, and 100mg/kg) on both locomotion and prepulse inhibition was assessed. The reanalysis incorporates data from 3 distinct animal groups administered with mescaline at doses of 10, 20, and 100mg/kg,

In this current study, an additional 17 groups of animals were subjected to the same experimental conditions as in the study of Páleníček et al., 2008 [19], including factors such as experimental setting, daytime, and season. These newly included groups encompass:

- Control group receiving a vehicle.
- Administration of 5HT1a, 5HT2a, 5HT2b, and 5HT2c antagonists individually.
- Co-administration of 5HT1a, 5HT2a, 5HT2b, and 5HT2c antagonists with 10, 20, and 100 mg/kg mescaline.

Subsequently, data from all 20 groups of animals were analyzed together.

### 2.1 Animals

A total of 410 experimentally naive Wistar male rats (SPF animals, 3-month old and obtained from Biotest Inc., Konárovice, Czech Republic) were housed in pairs in plastic cages and maintained in a temperature and light-controlled environment. The rats were provided with a standardized diet and water ad libitum. All experiments were conducted between 7 a.m. and 1 p.m. The study consisted of 20 experimental groups of animals, categorized according to the dose of mescaline and the specific antagonist administered. Each experimental group consisted of 9 to 10 animals for locomotor activity measurements (N=190) and 10 to 12 animals for sensorimotor gating testing (N=220). Each experimental subject was tested only once. The National Committee for the Care and Use of Laboratory Animals approved all experimental procedures, which were performed in accordance with the Animal Protection Act of the Czech Republic and respected the Guidelines of the European Union Council (86/609/EU).

### 2.2 Drugs and chemicals

To prepare for measurements, Mescaline.HCl (Sigma Aldrich) was dissolved in physiological saline and administered at doses of 10, 20, and 100 mg/kg, 60 minutes before testing. The selected serotonin receptor antagonists were also dissolved in physiological saline, and if applied, they were administered 15 minutes before testing. The following antagonists were used:

- The 5-HT1A antagonist N-[2-[4-(2-Methoxyphenyl)-1-piperazinyl]ethyl]-N-2-pyridinylcyclohexanecarboxamide maleate (WAY100635; Tocris Bioscience) at a dose of 1.0 mg/kg.
- The 5-HT2A antagonist (R)-(+)-α-(2,3-Dimethoxyphenyl)-1-[2-(4-fluorophenyl)ethyl]-4-piperinemethanol (M100907; Sigma Aldrich) at a dose of 0.5 mg/kg.
- The 5HT2B antagonist 6-Chloro-2,3-dihydro-5-methyl-N-5-quinolinyl-1H-indole-1-carboxamide (SB215505; Tocris Bioscience) at a dose of 1.0 mg/kg.
- The 5-HT2C antagonist 6-Chloro-2,3-dihydro-5-methyl-N-[6-[(2-methyl-3-pyridinyl)oxy]-3-pyridinyl]-1H-indole-1-carboxamide dihydrochloride (SB242084; Sigma Aldrich) at a dose of 1.0 mg/kg.

Control animals received a saline solution. All drugs were administered subcutaneously at a volume of 2 ml/kg.

### 2.3 Behavioral experiments

#### 2.3.1 Locomotor activity

In our previous publications [20–23], we provided detailed descriptions of the behavioral tests used, including both locomotion and PPI ASR. Briefly, locomotion (trajectory length) and its spatial characteristics (time spent in the center of the arena), were measured using a square black plastic box arena (68×68×30 cm) located in a quiet and evenly-lit room. Video recordings from a camera mounted above the arena were analyzed using EthoVision version 3.1.1 software (Noldus, Netherlands). At the beginning of each experiment, rats were placed in the center of the testing arena, and locomotion was measured for 30 minutes. To avoid the influence of odors from previous animals, the open field was cleaned with 10% ethanol and dried with a dry cloth after each test.

To determine the time profile after administration of mescaline and antagonists, locomotor activity was measured in 5-minute intervals and calculated by the EthoVision software. The time spent in the center of the arena (Tcenter) was evaluated using the EthoVision software by virtually dividing the arena into 5 × 5 zones, with 16 zones located peripherally and 9 centrally. The time spent in the center was calculated as the summation of time spent in the 9 central zones (Σt_1–9_; [24]).

#### 2.3.2 PPI ASR testing

The PPI ASR test was conducted using two startle chambers (SR-LAB, San Diego Instruments, San Diego, CA, USA) designed with a Plexiglas cylinder of 8.2 cm diameter and a piezoelectric accelerometer unit attached to a 12.5 × 3 × 25.5 cm Plexiglas platform. The background noise of 65 dB and the white noise acoustic stimuli (120-dB, 20-ms pulses and 70-dB, 20-ms prepulses) were controlled by the SR-LAB microcomputer and presented through high-frequency loudspeakers mounted above the cylinder. The piezoelectric accelerometer detected and transduced any vibrations caused by the whole-body startle response of the animal. The signals were digitized and recorded by a compatible computer interfaced with a startle apparatus. Sound levels were measured and calibrated with a Radio Shack Digital Sound Level Meter placed within each chamber. The experimental design was adapted from [25]. After a 5-minute acclimatization period with 65 dB background white noise, each rat was presented with five initial startle trials of 120 dB, which were not included in the analysis. Following these trials, each subject was exposed to four different types of trials presented in a pseudo-random order: 1/ single pulse (120 dB broadband burst, 20 ms duration), 2/ prepulse (13 dB, 20 ms duration above the background noise, presented 100 ms before the onset of pulse alone), 3/ prepulse alone (13 dB, 20 ms duration above the background noise) and 4/ no stimulus. Animals with a mean value lower than 10 were excluded from the calculation of the PPI ASR and were considered non-responders. The number of non-responders did not differ significantly between treatment groups.

#### 2.3.3 Statistical analysis

Statistical analyses were performed using Statistica v.13.3 (StaSoft Inc., Tulsa, Oklahoma, USA). Animals with z-scores more than 2 standard deviations from the mean were considered outliers and excluded from the analyses. To compare the effects of antagonists on mescaline-induced behavioral changes, a two-way analysis of variances (ANOVA) was conducted. Furthermore, a two-way repeated measures analysis of variances (RM ANOVA) was used to evaluate the effect of each antagonist on mescaline treatment, with treatment as the between-subjects factor and time interval as a repeated measures factor.

Where appropriate, the differences between groups were compared using Fisher’s LSD post hoc test with consecutive False Discovery Rate correction of individual P values. The alpha level of significance was set to p < 0.05.

## 3 RESULTS

### 3.1 Open field experiment

#### 3.1.1 Total locomotion

The results of the two-way ANOVA indicated significant main effects of dose (F(_3,176_) = 55.70; p < 0.001), treatment (F(_4,176_) = 30.77; p < 0.001), and the interaction between dose and treatment (F(12,176) = 10.37; p < 0.001). Post-hoc analysis revealed that the administration of mescaline alone at 100 mg/kg significantly increased trajectory length compared to saline, mescaline at 10 mg/kg, and mescaline at 20 mg/kg (p < 0.001).

Further comparisons of antagonist effects on mescaline-induced behavioral changes showed that M100907 effectively blocked the hyperlocomotion induced by mescaline at 100 mg/kg (p < 0.001). In addition, both M100907 alone and in combination with all doses of mescaline significantly decreased total locomotion compared to saline (p < 0.05-0.001). However, other antagonists administered alone or with lower mescaline doses (10 and 20 mg/kg) did not significantly change locomotor activity, but the difference between mescaline treatments and saline disappeared.

All remaining antagonists significantly reduced hyperlocomotion induced by mescaline at 100 mg/kg, with SB215505 showing the highest efficacy, followed by WAY100635 and SB242084 (p < 0.05-0.001). Only SB215505 was able to normalize the effects of mescaline to the level of saline animals, whereas the hyperlocomotion persisted to some extent following the administration of the other two antagonists (Figure 1).

**Figure 1.**
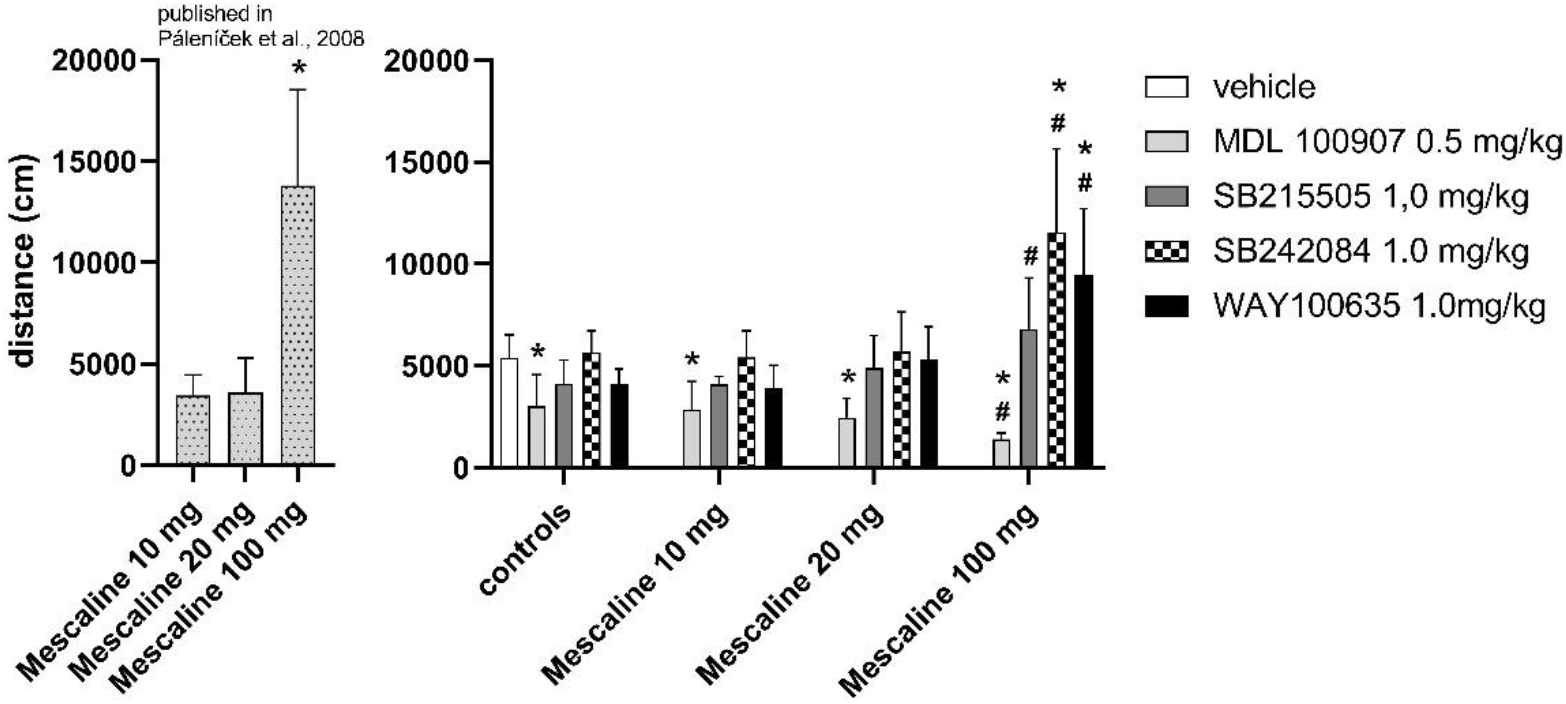
Open Field Test: Total locomotion - distance measured in cm over 30 minutes. Data are presented as means ± SD. Data that survived FDR correction are labeled as significant with asterisks indicating a significant differences from control (*; p < 0.05) and hashes a significant differences of groups treated with antagonists within each respective mescaline 10, 20 or 100 mg/kg alone treatment (#; p < 0.05).

#### 3.2.1 Time spent in the center

The results of the two-way ANOVA indicated that the treatment (F_(4, 174)_ = 10.46; p < 0.001), dose (F_(3, 174)_ = 4.06; p = 0.0081), and the interaction of treatment × dose (F_(12, 174)_ = 2.77; p = 0.0018) had significant main effects on the time spent in the center. Further post hoc analysis revealed that the highest dose of mescaline (100 mg/kg) significantly increased the time spent in the center when compared to the control vehicle and low doses of mescaline (p < 0.01 - 0.05).

Moreover, administration of all antagonists, except WAY100635, significantly attenuated the increased time in the center induced by mescaline at the highest dose (100 mg/kg) (p < 0.01 - 0.001; Figure 2), with the most pronounced effect observed with SB215505.

**Figure 2.**
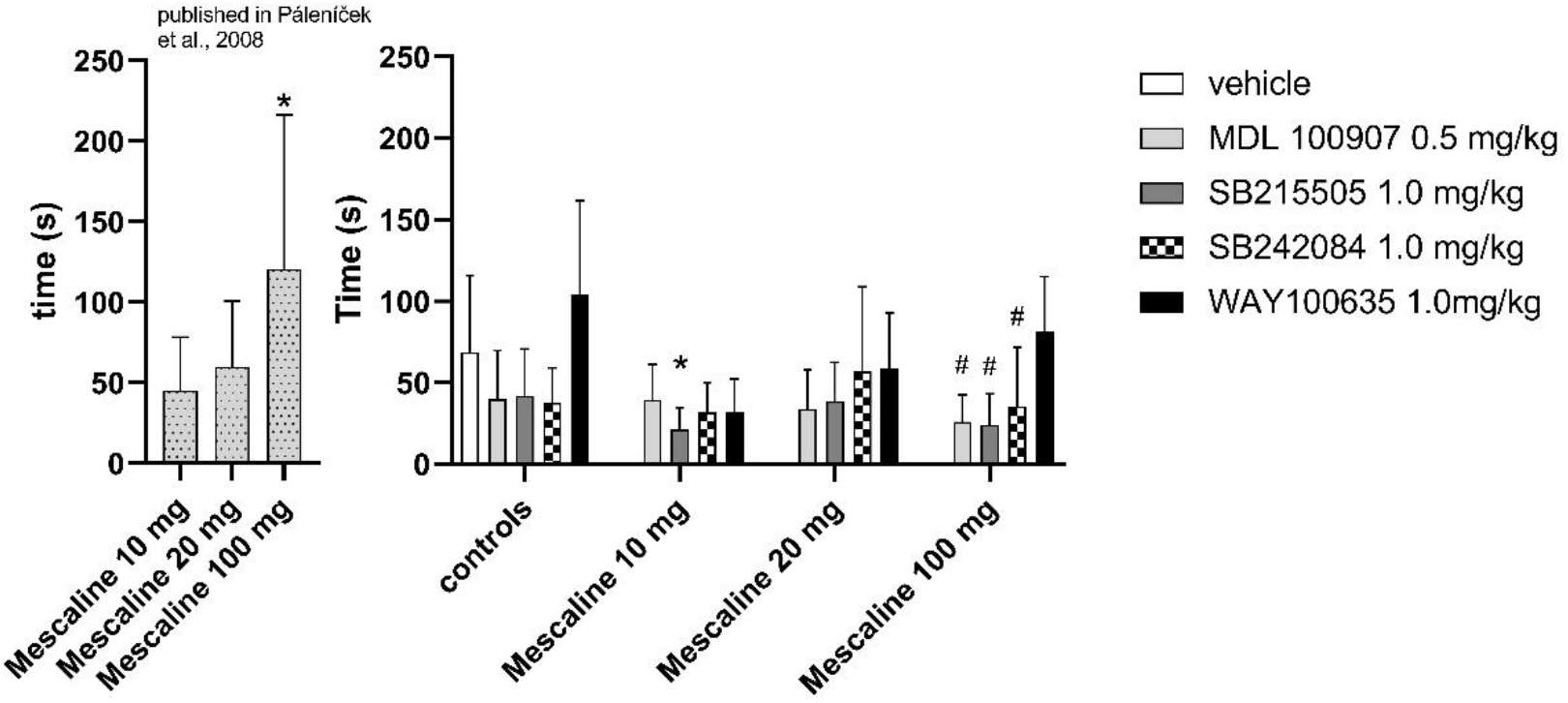
Open Field Test: Time Spent in Center – time in seconds. Data are presented as means ± SD. Data that survived FDR correction are labeled as significant with asterisks indicating a significant differences from control (*; p < 0.05) and hashes a significant differences of groups treated with antagonists within each respective mescaline 10, 20 or 100 mg/kg alone treatment (#; p < 0.05).

#### 3.1.3 Detailed analysis of 5 min intervals

A two-way RM ANOVA was conducted, revealing a significant main effect of treatment (F_(15, 143)_ = 23.11, p < 0.001), time (F_(5, 715)_= 393.83, p < 0.001), and a significant interaction between treatment and time (F_(75, 715)_ = 4.18, p < 0.001). Subsequent post hoc analyses demonstrated that low doses of mescaline (10 and 20 mg/kg) significantly decreased locomotion in the first (0-5 min) (p < 0.05) and second (5-10 min) time intervals (p < 0.05), while mescaline 100 mg/kg significantly increased locomotion compared to saline in all time intervals (p < 0.05 – 0.01). Administration of all antagonists, except M100907, partially normalized the hypolocomotion induced by mescaline 10 and 20 mg/kg within the first 5 minutes (p < 0.01 – 0.001). In contrast, only M100907 led to a significant blockade of hyperlocomotion induced by mescaline 100 mg/kg in all time intervals (p < 0.05 – 0.001) (see Figure 3).

**Figure 3.**
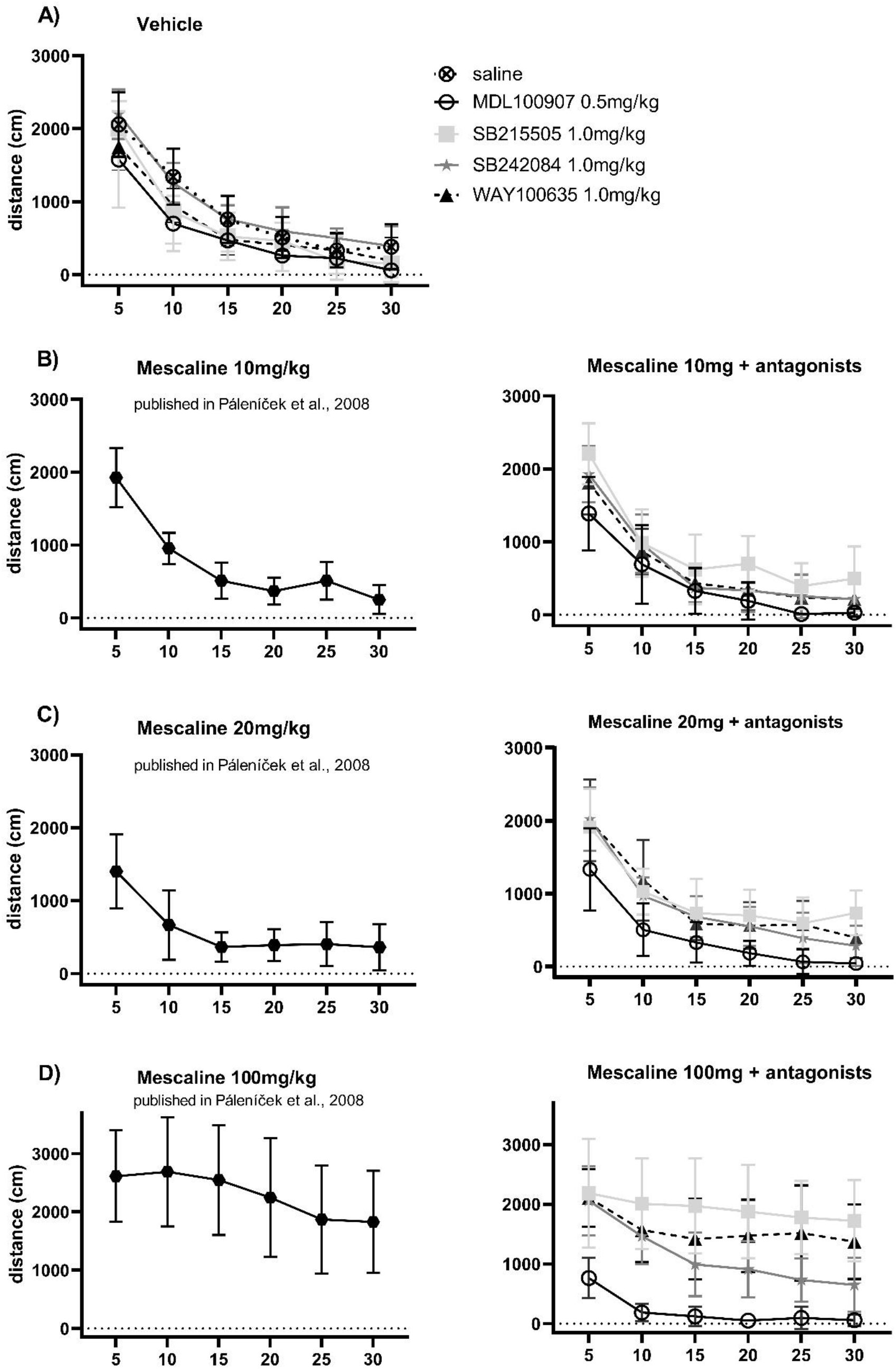
Open Field Test: 5-min intervals. Trajectory length measured in 5-minute intervals (distance in cm). Data are presented as means ± SD. **A)** Locomotion after administration of vehicle and antagonists alone. **B)** Locomotion after administration of mescaline at 10 mg/kg alone (left) and mescaline at 10 mg/kg with antagonists (right). **C)** Locomotion after administration of mescaline at 20 mg/kg alone (left) and mescaline at 20 mg/kg with antagonists (right). **D)** Locomotion after administration of mescaline at 100 mg/kg alone (left) and mescaline at 100 mg/kg with antagonists (right).

### 3.2 Prepulse inhibition of acoustic startle reaction (PPI ASR)

The results of a two-way ANOVA showed that there was a significant main effect of dose (F _(3, 182)_ = 5.038; p = 0.022) and treatment (F _(4, 182)_ = 13.333; p < 0.001) on ASR. However, post hoc analysis did not reveal a significant effect of mescaline treatment on ASR alone (refer to Table 1 for results).

**Table 1.**
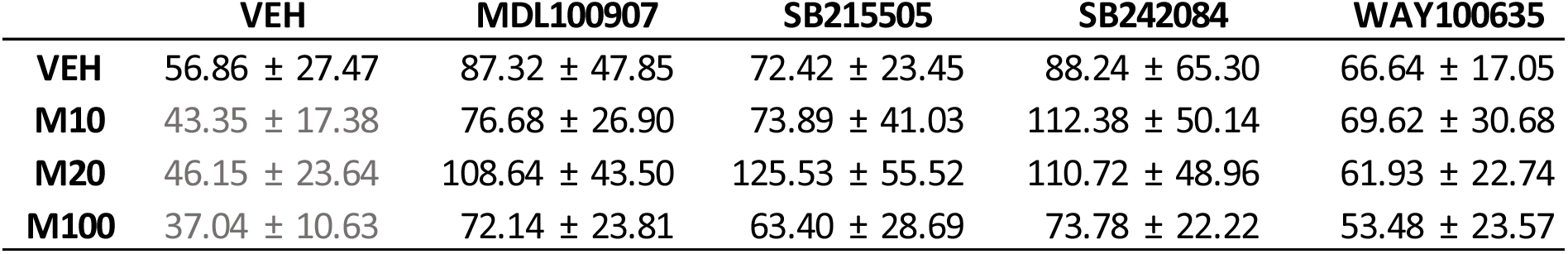
Acustic Startle Reaction. Amplitude in arbitrary units. Data published in Páleníček et al., 2008 are highlighted in grey. Data are presented as means ± SD.

Regarding PPI ASR, the two-way ANOVA indicated the main effects of treatment (F _(4, 182)_ = 5.960, p < 0.001) and dose (F _(3, 182)_ = 9.971; p < 0.001). Further analysis revealed that only M100907 restored PPI ASR deficit for all mescaline doses (p < 0.05 - 0.01) and SB242084 at a dose of 20 mg/kg of mescaline (p = 0.0017; refer to Figure 4).

**Figure 4.**
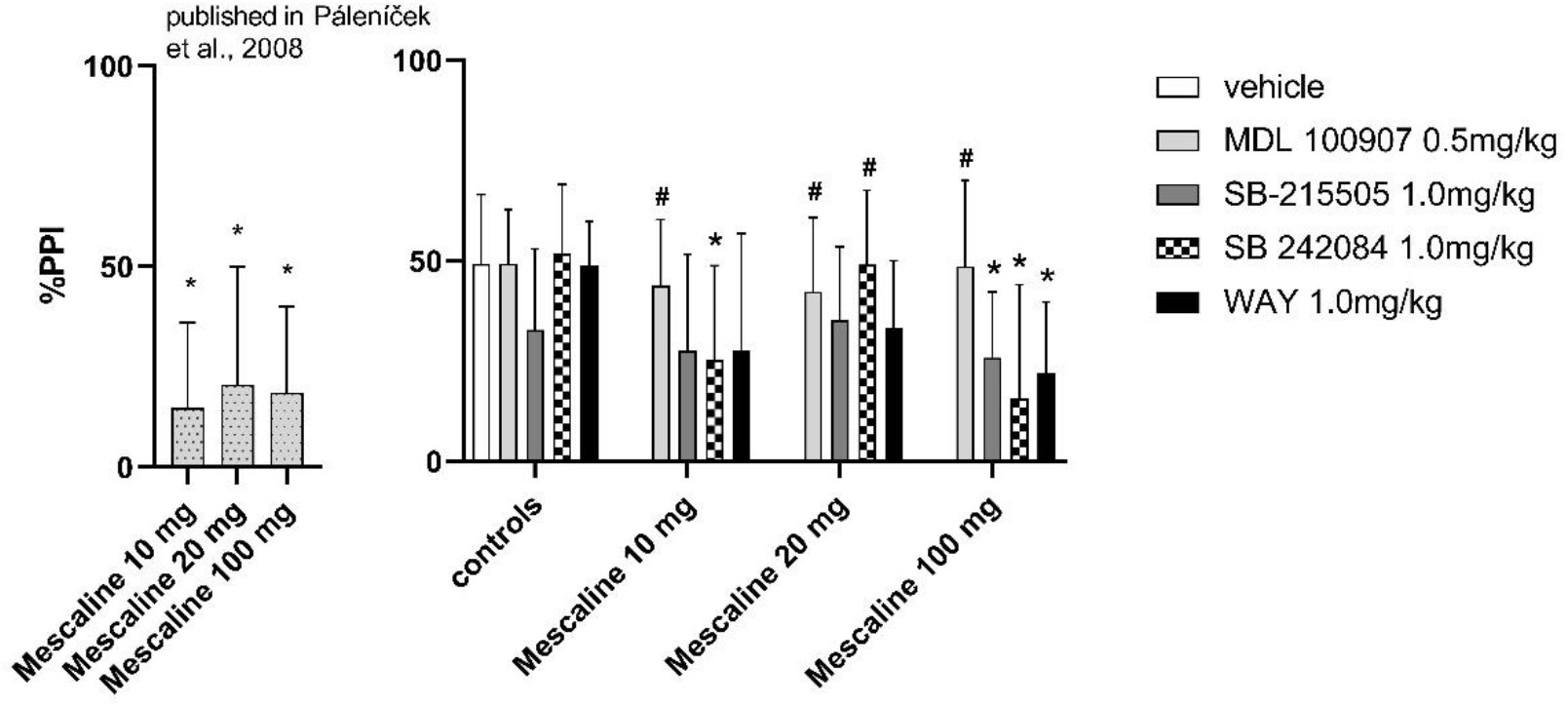
Prepulse Inhibition (PPI) of Acustic Startle Reaction (ASR). Percentage of PPI of ASR (%).Data are presented as means ± SD. Data that survived FDR correction are labeled as significant with asterisks indicating a significant differences from control (*; p < 0.05) and hashes a significant differences of groups treated with antagonists within each respective mescaline 10, 20 or 100 mg/kg alone treatment (#; p < 0.05).

## 4 DISCUSSION

### 4.1 Locomotor activity

The trend of decreased locomotion after mescaline administration as well as its biphasic action is in line with other behavioral studies in rodents [26]. The findings that 5-HT2A and 5-HT2C antagonist has an opposite role in the control of locomotion has been shown also in our previous psilocybin study in rats [12] and similar finding following serotonergic psychedelic DOI has been also described in mice [15]. As the crucial neurotransmitter involved in motor control is dopamine [27] it is highly probable that the observed effects are linked to the indirect modulation of dopaminergic tone by the 5-HT2A/C challenge. Indeed, whereas stimulation of 5-HT2A receptors induces a phasic increase in the release of dopamine, stimulation of 5-HT2C receptors decreases the tonic activity of the dopaminergic system [28,29]. Mescaline, which potently agonizes 5-HT2C receptors [3], would therefore decrease the tonus of the dopaminergic system resulting in hypolocomotion and a 5-HT2C antagonist, in turn, would increase basal dopaminergic activity, resulting in an improvement of locomotor deficit. Potent 5-HT2C antagonist SB242084 had been proven to have over 100-fold selectivity over a range of other 5-HT, dopamine, and adrenergic receptors [30], and also exhibited a potentiated effect on the locomotor activity stimulated by cocaine administration [31]. On the other hand, the complete reversal of hyperlocomotion following the highest dose of mescaline used by a 5-HT2A receptor antagonist supports the role of robust 5-HT2A dependent potentiation of phasic dopaminergic release that overcomes the inhibition related to decreased tonus mediated via 5-HT2C. The effects of 5-HT2C dependent inhibition of dopaminergic tonus are then unmasked by 5-HT2A antagonist resulting in almost complete suppression of locomotion.

Of interest are the effects of 5-HT2B and 5-HT1A antagonists which tended to normalize the mescaline-induced locomotor effects of both directions. The role of 5-HT1A receptors is not easy to study because they have opposite roles in terms of presynaptic and postsynaptic stimulation in the regulation of many behaviors including locomotion [32]. Interestingly in a study with cocaine, pretreatment with WAY100635 blocked the locomotor stimulant effect of cocaine which is known to be dopamine-mediated. Thus, regarding the inhibition of hyperlocomotor effects following the highest mescaline dose, similar mechanisms could be expected [33]. On the other hand, early pharmacological examination of WAY100635 demonstrated that it is not only a selective antagonist of 5-HT1A receptors, but it also possesses a lower affinity to α1-adrenergic and dopamine D2, D3 and D4 receptors [34,35]. Later, Chemel et al. [36] noticeably reported a relatively high affinity of WAY100635 to D4 receptors (16nM; 5-HT1A affinity is 2.2 nM) and confirmed low affinities at dopamine D2 and D3 receptors (940 and 370 nM, respectively). Moreover, they also perform functional assays in which it has been demonstrated that this compound is a potent agonist of the dopamine D4-mediated signaling pathway. Dopamine receptors agonism of WAY100635 could theoretically explain the partial efficacy in improving locomotor deficits induced by lower doses of mescaline.

The role of the 5-HT2B receptor in locomotor activity is not completely clear. Even though it had been demonstrated that the administration of SB-215505 caused increased locomotor activity and wakefulness in examined animals [37], the study of Reavill et al. [38] showed that SB215505 did not block the haloperidol-induced catalepsy and, also, SB-215505 did not alter cocaine-induced locomotor activity in the study of Fletcher et al. [31]. Also, the affinition of SB215505 to dopaminergic receptors was not examined yet and it most likely acts only through the blockade of 5-HT2B receptors. 5-HT2B receptor agonist, BW723C86, had shown a potent anxiolytic effect in several behavioral studies which were reversed by SB215505 [39,40], so the effect of the 5-HT2B antagonist on locomotor activity could be possibly caused by its anxiogenic effect.

### 4.2 Time spent in center

The increased time spent in the central parts of the arena following the highest mescaline dose (100 mg/kg) could be attributed to two plausible hypotheses. It may be directly linked to a general elevation in locomotor activity, or alternatively, the heightened presence in the central areas might signify a reduction in anxiety. It is also conceivable that this behavior is a result of a combination of both factors. The latter mentioned however seems contradictory to other studies with 5-HT2A agonists which seem to have rather anxiogenic-like effects [41,42]. As there was a non-specific effect of all antagonists it is likely supportive for the association with overall stimulatory activity. The role of anxiogenic/anxiolytic effects of tested compounds could however contribute to some extent as well. The anxiogenic-like effect of 5-HT2B antagonist SB-215505 was already discussed in subsection 4.1 and explains the decreased time in the center after its administration. The 5-HT2A activation is known to promote or increase anxiety-like behavior, whereas the 5-HT1A activation inhibits anxiety-like behavior [43,44]. Moreover, WAY100635 alone significantly increased anxiety-like behavior in an animal model of PTSD, while after inhibition of 5-HT2AR using ketanserin, the mice showed no statistical difference in most behavioral tests compared with the PTSD group [45]. Reported anxiogenic-like effects of 5-HT1A and 5-HT2B antagonists are in line with the results of our study. In contrary with our study, non-selective 5HT2A/2C receptor antagonists ritanserin and ketanserin administration are known to produce anxiolytic-like effects in rats during testing in the EPM [46,47]. Also, antagonizing 5-HT2A receptors due to M100907 administration was found to block methamphetamine-induced anxiety-like behaviors in rats [48]. As the 5-HT2A antagonist in our case resulted in almost complete immobility following mescaline treatment, it would be very difficult to evaluate its potential contribution in an anxiolytic/anxiogenic manner.

### 4.3 Sensorimotor gating

The PPI deficit was effectively normalized only by pretreatment with a 5-HT2A antagonist, so it depicts the most probable mechanism being related to 5-HT2A agonism. This is congruent with other studies with both indoleamine [e.g., LSD] [49] and phenylethylamine (e.g., DOI) hallucinogens in rats [16,50,51]. A tendency to normalize the PPI deficit induced by psilocin by 5-HT2A antagonist was congruently shown in our previous study [12].

### 4.4 Hypothetic non-serotonergic mechanisms

Finally, it should be noted that mescaline interacts with other receptors in the CNS such as alpha-adrenergic 2 receptors (Ki 1.4 µM) and it is thus possible that these receptors may contribute to the modulation of behavioral effects. It has been found that the stimulation of alpha1b- and alpha2-ARs increases DA-mediated locomotor response in both rats and mice [52]. Furthermore, mescaline has been shown to bind with some affinity (Ki 3.3 µM) to trace amine-associated receptor 1 (TAAR1) [53]. TAAR1 is also expressed within mesolimbic DA pathways [54,55] and interacts with both the DAT and the D2R [56,57] and the TAAR1 agonism was reported to reduce monoamine stimulation [58]. Since the dopaminergic transmission is crucial in motor activity and could be affected by the activity of both - adrenergic receptors and TAAR1, both receptors could contribute to the effect of mescaline in behavioral tasks. Therefore, serotonergic as well as non-serotonergic receptors might also contribute to the effects of mescaline in either effective or non-effective clinical doses.

## 5 CONCLUSION

The primary findings of the study indicate that the biphasic effects of mescaline on locomotion are regulated in opposite directions by two serotonin receptors, specifically 5-HT2A and 5-HT2B/C receptors. The initial ataxia is neutralized by 5-HT2B/C antagonists, while the hyperlocomotion is suppressed by 5-HT2A antagonism. The 5-HT1A antagonist partially normalized both effects. Moreover, the PPI deficit was found to be mediated by 5-HT2A-dependent mechanisms, while other receptor systems had only minor effects.

## 8 AUTHORS CONTRIBUTION

JH and TP were responsible for the study concept and design. LO-L drafted the manuscript. LO-L, MF-L and HD contributed to the acquisition, analysis, and interpretation of behavioral data. KŠ and KS assisted with data analysis and interpretation of the findings. TP and JH provided critical revision of the manuscript for important intellectual content. All the authors made a significant contribution to this study, read, revised, and gave final approval for the current version of the work to be published.

